# Cross-species permissivity: structure of a goat adeno-associated virus and its complex with the human receptor, AAVR

**DOI:** 10.1101/2022.01.14.476406

**Authors:** Edward E. Large, Mark A. Silveria, Tommi A. White, Michael S. Chapman

## Abstract

Adeno-associated virus (AAV) is a small ssDNA satellite virus of high interest (in recombinant form) as a safe and effective gene therapy vector. AAV’s human cell entry receptor (AAVR) contains Polycystic Kidney Disease (PKD) domains bound by AAV. Seeking understanding of the spectrum of interactions, goat AAVGo.1 is investigated, because its host is the species most distant from human with reciprocal cross-species cell susceptibility. The structure of AAVGo.1, solved by cryo-EM to 2.9 Å resolution, is most similar to AAV5. Through ELISA studies, it is shown that AAVGo.1 binds to human AAVR (huAAVR) more strongly than do AAV2 or AAV5, and that it joins AAV5 in a class that binds exclusively to PKD domain 1 (PKD1), in contrast to other AAVs that interact primarily with PKD2. The AAVGo.1 cryo-EM structure of a complex with a PKD12 fragment of huAAVR at 2.4 Å resolution shows PKD1 bound with minimal change in virus structure, except for disordering of a neighboring surface loop. Only 4 of the 42 capsid protein sequence differences between AAVGo.1 and AAV5 occur at the PKD1 binding interface. These result in only minor conformational changes in AAVR, including a near rigid domain rotation with maximal displacement of the receptor by ~1 Å. A picture emerges of two classes of AAV with completely different modes of binding to the same AAVR receptor, but within each class atomic interactions are mostly conserved.

**IMPORTANCE:** Adeno-Associated Virus (AAV) is a small ssDNA satellite parvovirus. As a recombinant vector with a protein shell encapsidating a transgene, recombinant AAV (rAAV) is a leading delivery vehicle for gene therapy with two FDA-approved treatments and 150 clinical trials for 30 diseases. The human entry receptor huAAVR has five PKD domains. To date, all serotypes, except AAV5, have interacted primarily with the second PKD domain, PKD2. Goat is the AAV host most distant from human with cross-species cell infectivity. AAVGo.1 is similar in structure to AAV5, the two forming a class with a distinct mode of receptor-binding. Within the two classes, binding interactions are mostly conserved, giving an indication of the latitude available in modulating delivery vectors.

## INTRODUCTION

Adeno-associated viruses (AAVs) are small single-stranded DNA viruses that can be repurposed into effective and safe gene therapy delivery vehicles (1). Primate AAV serotypes are the dominant choice for gene therapy. Several structures have been determined, starting with that of AAV2 by X-ray crystallography (2). AAV coding regions consist of two major open reading frames (ORFs): *rep* and *cap*, encoding functions needed in viral replication/DNA packaging and the capsid protein respectively (3). The Cap ORF encodes phenotypes relevant in tissue tropism and immune recognition (4, 5).

AAV2 was instrumental in the discovery and characterization of a human proteinaceous AAV receptor (AAVR) (6) (Fig. 1B). AAVR is a glycoprotein (N-linked/O-linked) and contains three major protein regions. The N-terminus contains a <u>M</u>otif <u>A</u>t <u>N</u>-terminus with <u>E</u>ight-<u>C</u>ysteines (MANEC) while the central extracellular portion contains five Polycystic Kidney Disease (PKD) domains numbered 1 to 5 from N-terminus to C-terminus (7). The C-terminus has a predicted transmembrane protein domain and cytosolic domain responsible for trafficking AAV through the trans-Golgi network (TGN) (6). Experiments in mouse models and human cell lines indicate the physical interactions between primate AAVs and AAVR receptors of different species can be conserved.

**Figure 1:**
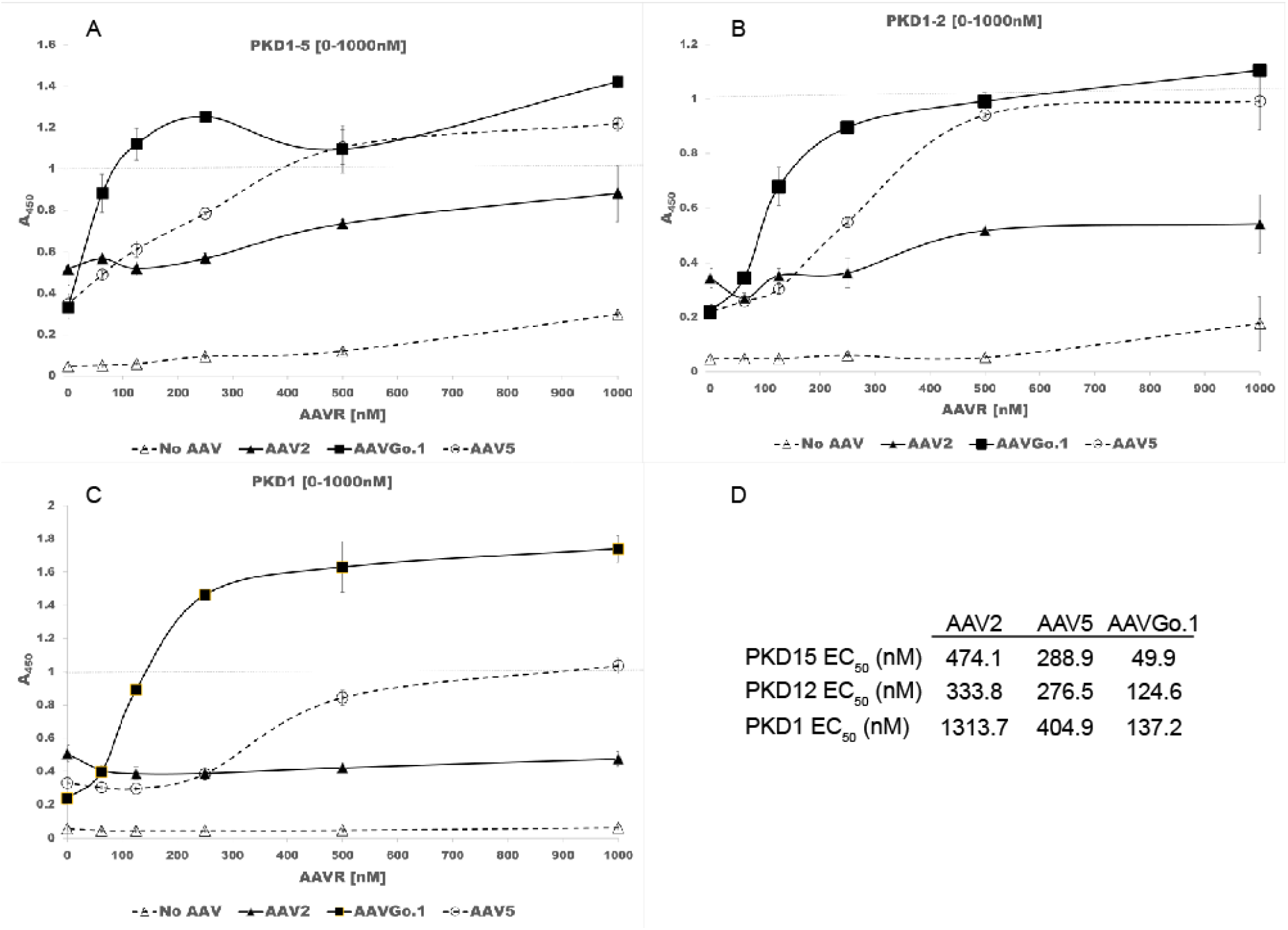
AAVGo. 1 interacts with PKD domains. A) PKD15 ELISA. B) PKD12 ELISA. C) PKD1 ELISA. D) EC_50_ values determined from panels A-C.

For human AAVR (huAAVR), structures for a handful of complexes are known (8–12). The structures consist of AAV1, AAV2 or AAV5 serotypes complexed with human PKD variants (huPKD) containing domains 1-2 (huPKD12) or domains 1-5 (huPKD15). Both structure and mutant data (13, 14) consistently indicate AAV5 is unlike all other AAVs whose interactions with huAAVR are mediated primarily through PKD2. Domain-swap mutants show AAV5 interacting exclusively with PKD1 (13) and the cryo-EM structures of complexes with huAAVR fragments show that the PKD1 binding site on AAV5 (9, 11) is different from the common PKD2 site for AAV2 and AAV1 (8, 9, 11).

Domain-swap mutants indicate a secondary role for PKD1 with AAV2 and other serotypes (13) but multiple cryo-EM structures reveal only the tightly-bound PKD2 at high resolution (8, 12, 15) It seemed possible that AAV2-like viruses bound not just to PKD2, but also to PKD1, though so weakly that PKD1 had not been seen in the AAV2 structures. It was not as simple as overlaying PKD1 in an AAV5-like position and connecting it, as a single subunit chain, to PKD2 as found in AAV2, because connecting the domains required implausible stereochemistry (11).

AAV gene therapy utilizes primate serotypes of which AAV5 is currently the only known representative with primarily PKD1 interactions. AAV5 was initially isolated from a human penile lesion (Bantel-Schaal and Zur Hausen 1984) and is the sole primate AAV5 clade representative. The AAV5 clade also includes AAVGo.1 (16, 17), which was isolated from deceased neonatal goat ileum (Olson, Haskell et al. 2004). Both recombinant AAV5 and AAVGo.1 can transduce at equivalent levels in primate and ruminant cell lines (Arbetman, Lochrie et al. 2005, Qiu, Cheng et al. 2006). Efficient transduction of primate and ruminant cell lines by both AAV5 and AAVGo.1 represents the broadest divergence between AAV host species that are reciprocally permissive to cross-species infection. Given the strong evidence for AAV5 interactions with PKD1 we investigated potential interactions of AAVGo.1 with huAAVR.

Here, binding of AAVGo.1, AAV5 and AAV2 to domain combinations of huAAVR are compared by ELISA. Then high-resolution structures are determined by cryo-EM of AAVGo.1 alone, and in complex with a PKD12 fragment of huAAVR. These are compared to corresponding structures of AAV5 to understand the diversity of interactions that are compatible with productive receptor-binding.

## RESULTS

### AAVGo.1 binds huAAVR

HuAAVR binds to AAV5, AAVGo.1 and AAV2 with comparable order-of-magnitude avidities as measured by ELISA binding using huAAVR ectodomain constructs. In contrast to AAV2, the binding of both AAVGo.1 and AAV5 is mediated mostly through the PKD1 domain. Both have stronger overall binding avidity than AAV2, AAVGo.1 having the strongest yet measured (Figure 1).

### Structure of AAV-Go.1 and its huAAVR receptor complex

Cryo-EM single particle analysis (SPA) yielded reconstructions at 2.9 Å resolution for native AAVGo.1 and 2.5 Å for the AAVR PKD12 complex. Sharpened maps provided detail for atomic modeling and were interpretable from residue 209 to 726 of the capsid protein in both reconstructions with one exception. In the receptor complex, residues 477 to 482 are less ordered and only traceable in maps low-pass filtered to 10 Å resolution. By convention, the numbering of VP1 is used, even though it constitutes only 10% of the capsid subunits. The majority of the subunit common to VP1-3 is seen in the 60-fold averaged reconstructions, starting at residue 209, before βA. The N-terminus of VP3 is residue 193 by VP1 numbering, so the atomic model accounts for 517 of 534 (96.8%) VP3 residues.

The native AAVGo.1 capsid has a similar protein fold and surface topology to AAV5 structures (9, 11, 18). The capsid core, comprised of a jellyroll fold found in many virus structures (19), and a single α-helix (αA) are the most conserved features of AAVs, and AAVGo.1 is no exception. The AAVGo.1 core jellyroll has the standard topology with backbone chain alternating between two stacked antiparallel β-sheets (almost rolling into a single tube). The sheet forming the inner surface of the capsid consists of β-strands B, I, D and G that alternate with the “CHEF” strands of the outer sheet, together with a 5^th^ inner strand, βA. βA is seen in many parvoviruses (20) at the edge of the BIDG sheet, running antiparallel to and turning directly into βB. Seven loops that connect strands of the BIDG and CHEF sheets encode much of the functionality of the capsid protein. Within the loops are regions of low sequence conservation that decorate the outer surface of the virus and these variable regions (VR) are enumerated as VR-I to VR-IX (21). VR-I (262-268) is found between βB and βC strands (BC loop), VR-II (362-330) is found in the DE loop, VR-III (380-388) is in the EF loop, VR-IV to VR-VIII (449-468, 487-504, 525-541, 544-556, and 579-594) are located in the GH loop and the C-terminus contains an additional VR-IX (704-711) after the last βI strand. These VR loops serve as guideposts for understanding AAV:AAVR interactions (8–12).

AAVGo.1 and AAV5 capsid proteins have high homology (94%). All 42 amino acid differences occurring in the VP3 region (more specifically C-terminal of AAVGo.1 VP1 residue 376) and the majority of these differences are located on the exterior of the capsid (17).

When comparing PKD1 as bound to AAVGo.1 or AAV5 we see an overall shift in the receptor position (Figure 3). Displacements of up to 1 Å are observed at some backbone carbons (Figure 3B, C). The shift appears to be a rotation about Arg_353_ which remains essentially unmoved (Figure 3D). Displacements become greater with distance from Arg_353_. What could cause this rotation between one serotype and another that are so similar? The key perhaps lies in the four residues that are changed in the binding footprint of AAVGo.1 when compared with AAV5.

The binding site formed by AAV5 and PKD1 has previously been examined (9, 11). Here, our attention is focused on the four viral residues within the binding interface that are different between AAVGo.1 and AAV5. Central to the binding site is human PKD1 residue Arg_353_, which makes contact with AAV5 Ser_531_ and Gly_545_. In AAVGo.1, the equivalent residues to Ser_531_ and Gly_545_ are, respectively, Ala_533_ and Asp_547_ (Figure 2C, D & Figure 4B, C). Both AAV5 Ser_531_ and AAVGo.1 Ala_533_ make contact (exclusively) with Arg_353_ (≤ 3.5 Å). AAVGo.1 Ala_533_ contacts AAVR Arg_353_ while the corresponding AAV5 Ser_531_ adds to the surface contact. Neighboring AAV5 Gly_545_ makes minimal contact with AAVR Arg_353_, while the corresponding AAVGo.1 Asp_547_ makes several van der Waals contacts in addition to AAVR Arg_353_ (Figure 4B). In AAVGo.1 Pro_545_ replaces AAV5 Leu_543_ (Figure 2E, F & Figure 4D). Both are involved in somewhat different virus-receptor van der Waals contacts, probably of similar extents. Lastly, in AAVGo.1 Ser_542_ replaces AAV5 Ala_540_, adding a hydroxyl group that forms a polar interaction with AAVR Asn_395_ (Figure 2E, F & Figure 4D). In summary, there are some detailed differences, the net effect of which is a modest increase in contacts and polar interactions for the AAVGo.1 complex compared to AAV5.

**Figure 2:**
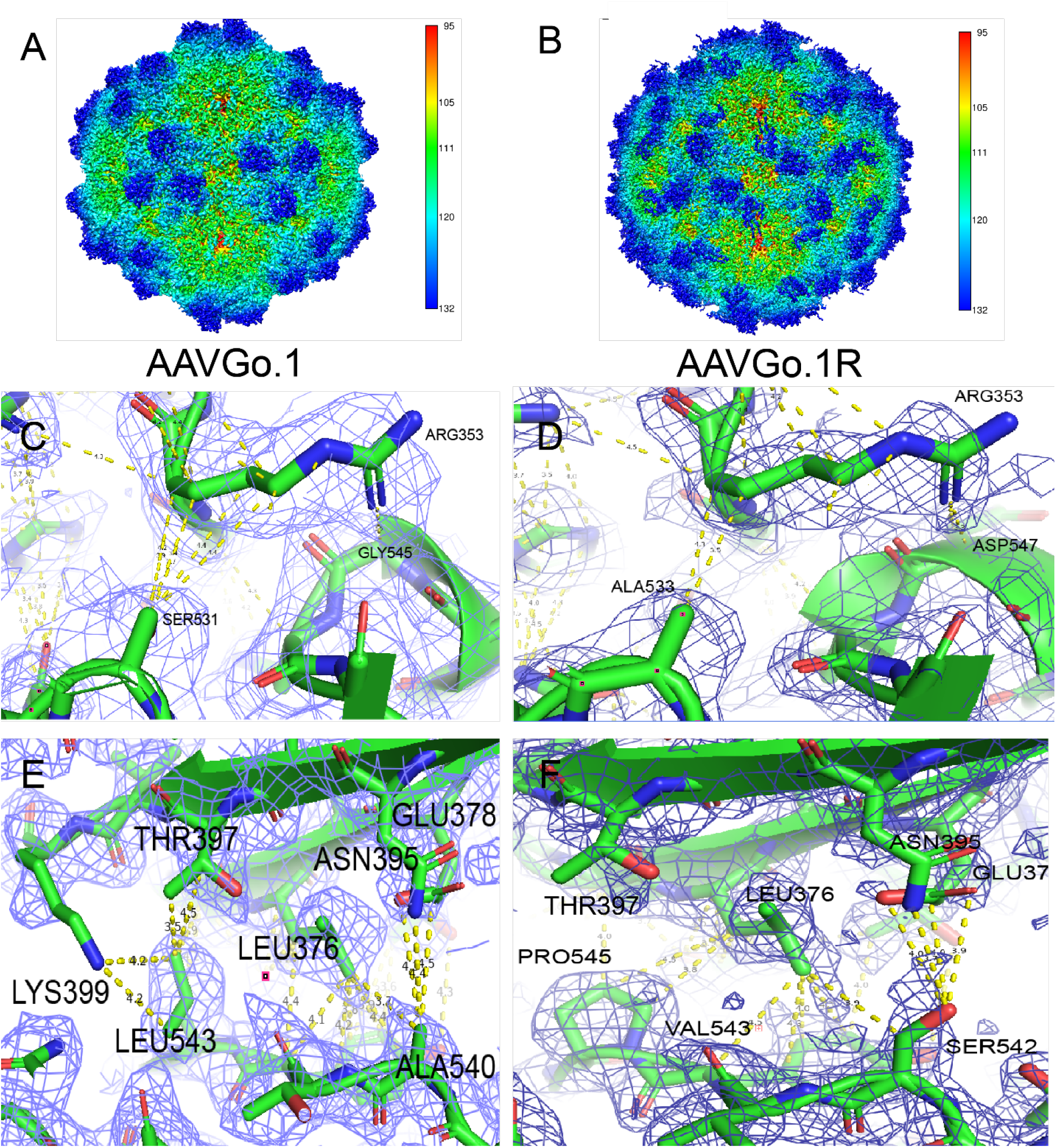
The receptor site on AAVGo is similar to AAV5 with some key differences. A) Native reconstruction of AAVGo. 1, colored by distance from the virus center. B) AAVGo. 1 complexed with PKD1. The AAV5 PKD1 N-terminus begins above the edge of the 5-fold pore and descends towards the 2-fold axis (Silveria, Large et al. 2020). Panels C – E together show the environment for the four interface residues that differ between AAV5 (left) and AAVGo. 1 (right). Atomic models are overlaid on maps contoured at 1.5 σ. C) The AAV5 environment around AAVR-Arg_353_, AAV5-Ser_531_ and AAV5-Gly_545_. D) The corresponding AAVGo. 1 environment around huAAVR-Arg_353_, AAVGo. 1-Ala_533_ and AAVGo. 1-Asp_547_. E) The AAV5 environment around Leu_543_ and Ala_540_. D) The corresponding AAVGo. 1 environment around Pro_545_ and Ser_542_.

**Figure 3:**
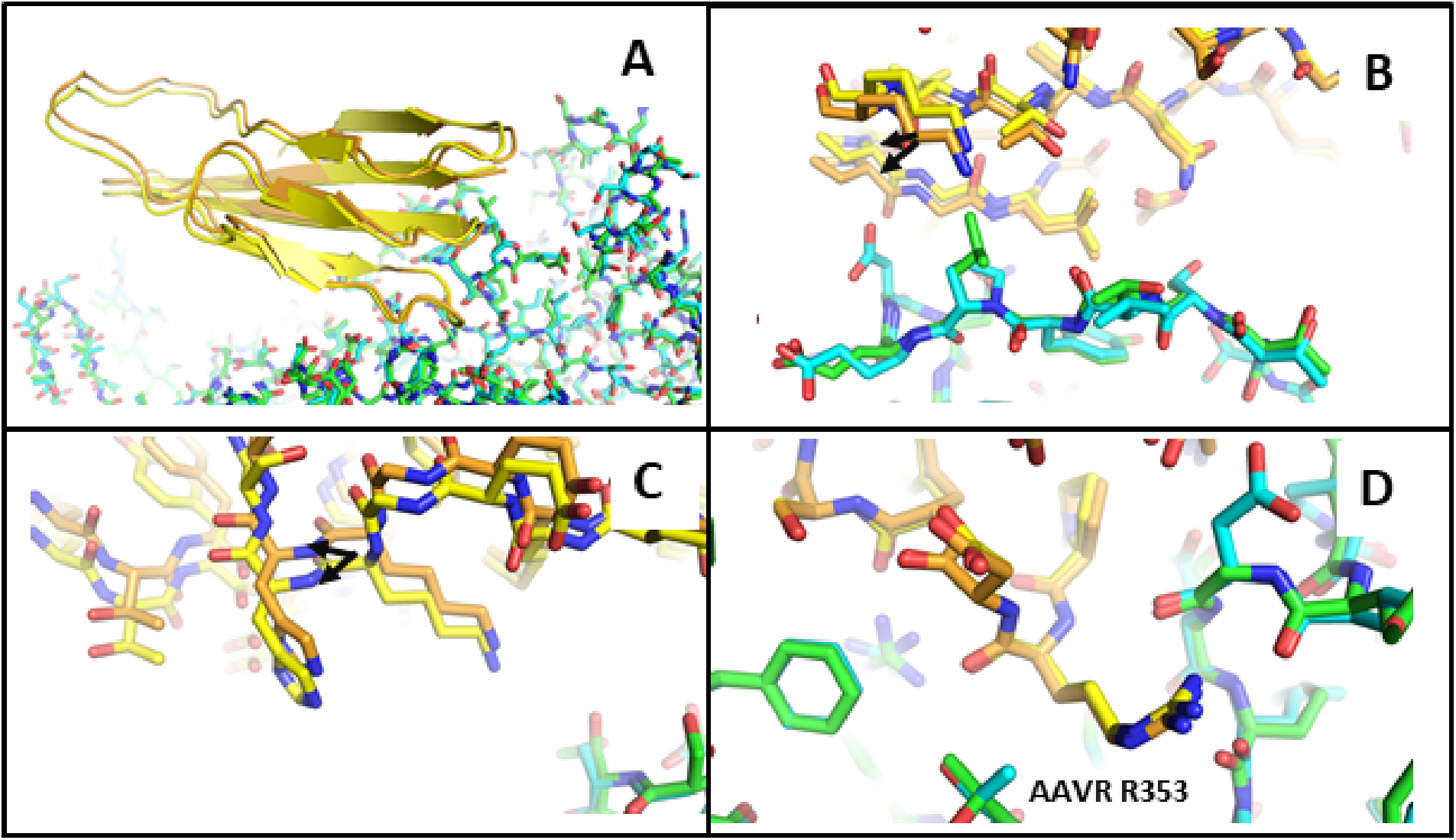
A rotation in the PKD1 position is observed when comparing its bound structure to AAVGo. 1 versus AAV5. AAVGo. 1 is colored with cyan carbons while AAV5 is colored with green carbons. The PKD1 from the AAVGo. 1 structure is colored with yellow carbons and the PKD1 from the AAV5 structure is colored with orange carbons. A) The overall rotation is observed in the PKD1 position when bound to either AAVGo. 1 or AAV5. B) A close up shows the PKD1-viral interface near AAVR Lys_399_. Arrows indicate a pair of corresponding atoms that differ by ~1 Å. C) This view is near AAVR His_363_. D) On the lowermost turn in panel A, AAVR Arg_353_ is bound more closely in a pocket on the AAV surface, and there is little difference between AAVGo. 1 and AAV5. Thus the overall displacement is approximately a rotation about an R353 pivot point.

**Figure 4:**
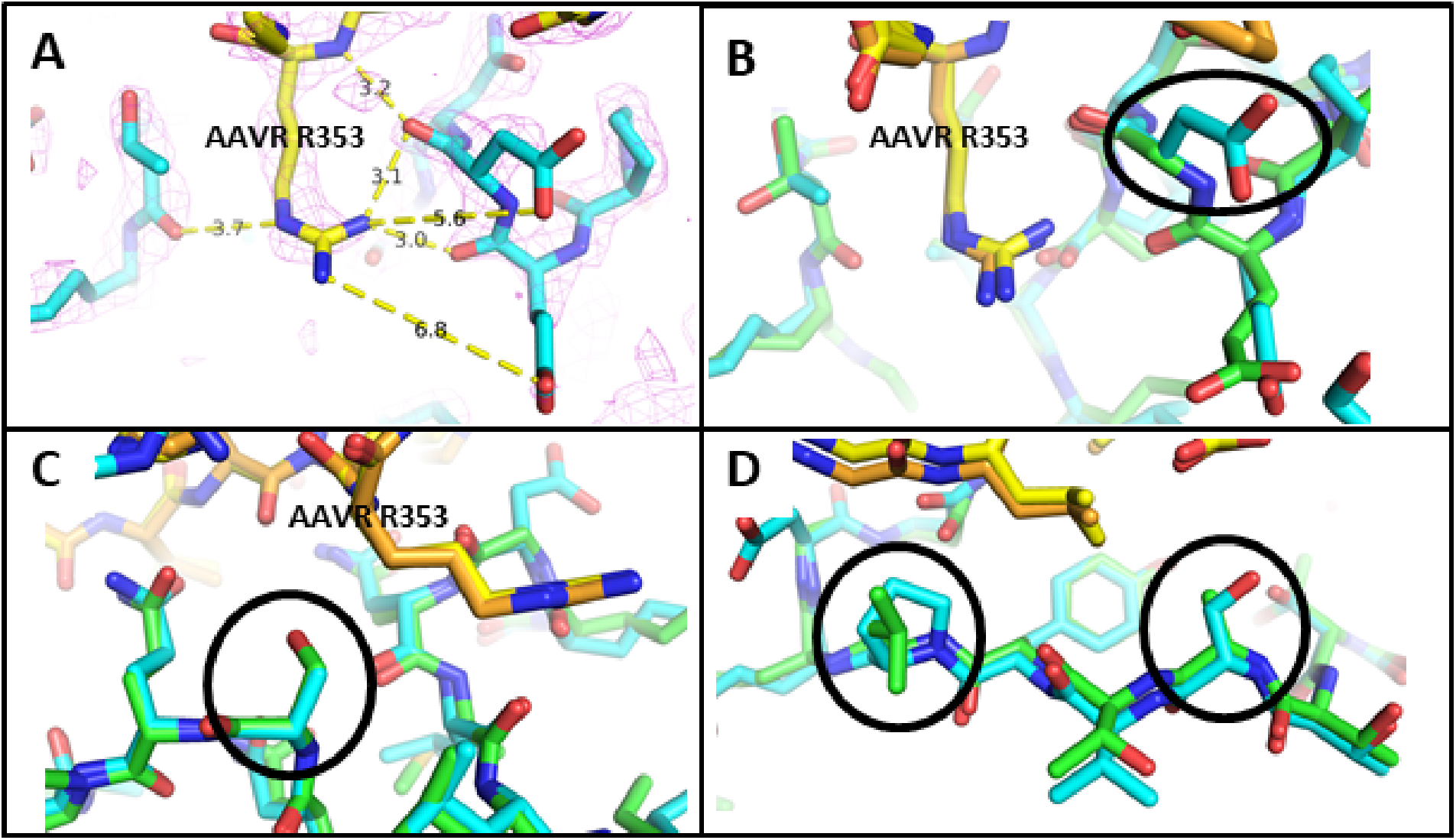
Comparison of the virus-receptor complexes of AAVGo. 1 with AAV5. A) AAVR (yellow carbons) bound to AAVGo. 1 (cyan carbons) near AAVR-Arg_353_. B) The AAV5 complex structure (PDBid 7kpn) is overlaid with green / orange carbon atoms. Circled are the substitutions of Asp_547_ for Gly_545_ in AAVGo. 1 (vs. AAV5) providing a more complementary, negatively charged binding pocket for AAVR-Arg_353_. C) Circled is the substitution of AAVGo.1 Ala_533_ for AAV5 Ser_531_. D) Within the circles, Pro_545_ and Asp_547_ replace AAV5 residues Leu_543_ and Gly_545_.

The largest difference, by sequence, is a double insertion in AAVGo.1 relative to the AAV5 sequence: L_477_(SS)G_478_. The region is disordered in AAVGo.1 receptor complex, lacking density between Thr_477_ and Ser_482_ in the sharpened map. When low-pass filtered to 10 Å resolution, the backbone can be traced readily, indicating that the region is dynamic. In unbound AAV5 and AAVGo.1, this loop is not disordered, and it appears less disordered in the AAV5-AAVR complex. Thus, it does appear that the combination of receptor-binding and the tandem serine insertion increases the disorder. Indeed, the map recovered by low-pass filtering is close to the C-terminal end of PKD1, discussed below.

## DISCUSSION

The largest difference in sequence between AAVGo.1 and AAV5 is the tandem serine insertion in variable region V (VR-V), a site that is more mobile than other regions of the surface, witness the fact that it is observed only in a lower resolution map low-pass filtered to 10 Å. The difference is within the epitope of a neutralizing monoclonal antibody, mAb HL2476 (22), and if the mAb were to be representative of natural neutralizing polyclonal antibodies, it is possible that the difference reflects selection through immune-evasion. We note that the loop that is extended in AAVGo.1 by the two-residue insertion is in the general vicinity of the final residue observed in PKD1, about 6 Å away, and one cannot exclude the possibility of an additional interaction between VR-V of AAVGo.1 and the interdomain linker between PKD1 and PKD2 in AAVR.

Prior to this work, most complexes between AAV and AAVR were of similar structure, like the AAV2 complex (8, 9, 12), with a single exception, that of AAV5 (9, 11). At first approximation, the binding of AAVGo.1 to huAAVR is similar to that of AAV5, so the single exception becomes a second class in terms of the mode of binding. Between the two classes, the modes of binding are completely different. Within each class, the diversity in structure of the virus-receptor interface is much more modest. It is anticipated that within and between binding classes, diversity will yield distinct transduction efficiencies, tissue tropism and antibody neutralization profiles.

## MATERIALS AND METHODS

### Expression and purification of AAVGo.1 and AAVR constructs

AAVGo.1, AAV2 and AAV5 Virus-Like Particles (VLPs) were produced using Sf9 cells and purified via ultracentrifugation in a cesium chloride gradient, repeated 4 times, and yielding a final concentration of 38.8 mg/ml (10). Sample purity and concentration were evaluated using nanodrop, SDS-PAGE and negatively stained electron microscopy using a JEOL JEM-1400 120kV Transmission Electron Microscope (TEM). AAVR fragment constructs (His_6_PKD1; His_6_PKD12; and His_6_PKD15) were expressed and purified as previously described (10).

### Enzyme-Linked Immunosorbent Assay (ELISA)

ELISA assays were carried out in triplicate using a modified direct ELISA protocol (6, 11). AAV2, AAV5, or AAVGo.1 VLPs (2.5 μg/ml) were incubated in ELISA plates (Corning Costar # 9018) in 100 mM NaHC03 (pH 9.6) and washed with TBST buffer (0.05% TWEEN in TBS). Plates were incubated with N-terminally His-tagged AAVR (His_6_PKD1; His_6_PKD12; or His_6_PKD15) and detected with anti-6x His tag horseradish peroxidase (HRP) (abcam #ab1187) antibody. 3,3’5,5’-Tetramethylbenzidine (TMB) ELISA Substrate (abcam #ab171523) was added to each well and development was stopped using 1 M hydrochloric acid. The optical density of plates was evaluated at 450 nm using a microplate reader (BioTek Synergy H1 Hybrid Multi-Mode Reader). EC_50_ values were estimated using the ATT BioQuest^®^ - EC50 calculator.

### TEM/Single Particle Cryo-EM data acquisition and 3D image reconstruction

Native AAVGo.1 and AAVGo1:PKD12 complexes were prepared and imaged as previously described for AAV2 and AAV5 (11, 12). Briefly, AAVGo.1 was dialyzed/diluted into HN buffer (25 mM HEPES pH 7.4 and 150 mM NaCl) at 0.33 mg/ml and PKD12 at 0.75 mg/ml. Grids had an ultrathin continuous carbon film layered on Lacey carbon supports (400 mesh; copper; Ted Pella # 01824). The grids were glow discharged using a PELCO easiGlow ™ Glow Discharge Cleaning System. After application of 2 μl of sample, native AAVGo.1 grids were blotted once and plunged directly into liquid ethane using an FEI Mark IV Vitrobot (force = 4, time = 2, temp = 25, humidity = 100). PKD12 complexes grids were prepared by adding 2 μl AAVGo.1 to the grid with a 2 minute incubation time. The grid was then wicked with Whatman filter paper (grade 595) and 2 μL of receptor was immediately added followed by blotting and plunging via Vitroblot (force = 4, time = 3, temp = 25, humidity = 100).

Cryo-EM grids were screened using an FEI Tecnai F30 Twin 300 kV TEM in preparation for single particle Cryo-EM. Native AAVGo.1 and AAVGo.1:PKD12 complexes were imaged using an FEI Titan Krios equipped with a Gatan K3 digital camera in super-resolution mode at a dose rate of 40 frames per 3.12 second exposure (Table 1). Automated micrograph data collection was enabled using Leginon (23).

**Table 1:** Cryo-EM data acquisition and processing

| **Data Collection** | **AAVGo.1** | **AAVGo.1R** |
| --- | --- | --- |
| Magnification (x) | 64,000 | 64,000 |
| Voltage (kV) | 300 | 300 |
| Electron Exposure (e/Å<sup>2</sup>) | 35.4 | 32.9 |
| Defocus Range (µm) | -0.7 to -2.0 | -0.7 to -2.0 |
| Pixel Size (Å) | 0.664 | 0.664 |
| **Data Processing** |  |  |
| Motion Correction | Relion 3.1.1<br>CTFFIND- | Relion 3.1.1 |
| CTF Estimation | 4.1 | CTFFIND-4.1 |
| Symmetry Imposed | I1 | I1 |
| Initial Particle Images | 246,094 | 208,921 |
| Final Particle Images | 140,568 | 71,095 |
| Map Resolution | 2.93 | 2.36 |
| FSC Threshold | 0.143 | 0.143 |

Relion 3.1.0 was used to process the data for native AAVGo.1 (24). 713 micrographs were motion corrected using Relion’s own implementation and CTFFind-4.1 was used for CTF correction (25). A total of 246,094 particles were picked with Relion autopick, which were culled down to 188,650 through several rounds of 2D classification. An additional round of 3D classification resulted in 140,568 particles. 3D refinement produced a map of 3.62 Å (gold-standard FSC_0.143_). Post-processing, including per particle CTF and motion corrections and masking resulted in a final resolution of 2.93 Å.

Relion 3.1.1 was used for the complex of AAVR PKD12 with AAVGo.1 (24). A total of 1204 micrographs were motion corrected with MOTIONCOR2 (26) and CTF estimation was carried out using CTFFind-4.1 (25). From 1127 micrographs, 208921 particles were picked using Relion’s Autopick with the native AAVGo.1 structure as a template. 3D Refinement was carried out directly on the extracted particles and 3D classification was used with C1 symmetry imposed resulting in 71095 particles for 3D reconstruction. Per particle CTF and motion correction were performed followed by 3D auto-refinement with I1 symmetry resulted in an unmasked structure at 2.55 Å. Masking resulted in a final resolution of 2.36 Å (gold-standard FSC_0.143_).

### Model building and structure refinement

A preliminary model for native AAVGo.1 was generated using the native AAVGo.1 VP3 protein sequence threaded into the 2.1 Å AAV5 structure (PDBid: 7kp3) in Coot (27). The starting model for the PKD1 domain of the receptor came from the 2.5 Å AAV5:PKD1-2 structure (PDBid: 7kpn). The models and imaging parameters were refined using RSRef 0.5.6 (28) with a final real-space correlation coefficient (CC) of 0.87 for native AAVGo.1 and 0.83 for complexed AAVGo.1:PKD1. (Correlation coefficients were calculated using all map grid points within 2.0 and 2.4 Å of all atoms, respectively.) Manual model adjustments were executed in Coot 0.8.9.2.

## ACKNOWLEDGMENTS

The Chapman lab is grateful for the contributions of the University of Missouri Electron Microscopy Core the Purdue Cryo-EM Facility. Thank you also to Dr. Jianming Qiu and Dr. David Pintel for AAVGo.1 and AAV5 Rep-Cap plasmids. This work supported by the National Institute of Health R35 GM122564. Purdue Cryo-EM Facility data acquisition was supported by National Institute of General Medical Sciences Grant U24 GM116789. Atomic models and cryo-EM maps will be made available at the PDB and EMDB.

